# Temporal trends in incidence of childhood cancer in Switzerland, 1985-2014

**DOI:** 10.1101/670224

**Authors:** Grit Sommer, Matthias Schindler, Shelagh Redmond, Verena Pfeiffer, Garyfallos Konstantinoudis, Roland A. Ammann, Marc Ansari, Heinz Hengartner, Gisela Michel, Claudia E. Kuehni, for the Swiss Paediatric Oncology Group (SPOG)

## Abstract

**Background:** Incidence of childhood cancer increased in most countries worldwide, but the reasons are unclear. This study investigates trends in childhood cancer incidence in Switzerland from 1985 to 2014.

**Methods:** We extracted data on all childhood cancer cases diagnosed at ages 0-14 years in Switzerland from the Swiss Childhood Cancer Registry. We included ICCC-3 main groups I-XII and calculated age-standardised, cumulative, and age-specific incidence for different diagnostic groups. We analysed trends in annual age-standardised incidence using JoinPoint regression models.

**Results:** Over the study period from 1985-2014, 5104 of 5486 cancer diagnoses (93%) were microscopically verified. The proportion of children treated in paediatric cancer centres increased from 84% during 1985-1994 to 93% in 1995-2004 and 98% in 2005-2014 (p<0.001). Using the 2010 European standard population, age-standardised incidence was 143 in 1985-1994, 154 in 1995-2004, and 162 per million in 2005-2014. Over the period 1985-2014, incidence for all cancers increased by 0.7% (95% confidence interval [CI] 0.5-1.0) per year, 0.8% (95% CI 0.2%-1.4%) for leukaemias, 3.8% (95% CI 1.7%-6.0%) for epithelial neoplasms and melanomas, and 3.0% (95% CI 1.3%-4.6%) for CNS tumours for the period 1985-2002.

**Conclusion:** Trends in incidence were driven mostly by increases among leukaemias and CNS tumours. For CNS tumours, observed trends may be explained at least partially by diagnostic changes and improved registration. For leukaemias, rising incidence may be real and at least partly due to changes in risk factors.

**Highlights:**

- In Switzerland, incidence of childhood cancer increased by 18% from 1985-2014.
- Increase in incidence was mainly caused by brain tumours and leukaemias.
- Improved registration and diagnostics may have increased brain tumour incidence.
- Increasing trend for leukaemias may be real, but reasons remain elusive.

## Introduction

Incidence of childhood cancer during the decade 2001-2010 increased worldwide by 13% compared to rates in the 1980s.^1^ The reasons for this increase remain unclear. Improved case ascertainment and diagnostics could account for this trend,^2,3^ or the trend could represent a real increase in cancer risks.^4-8^ Most countries have seen increasing trends in childhood cancers, though results vary by reporting period and cancer type.^2,8-20^ Switzerland is one of the few countries with a national childhood cancer registry that has been in existence for over 40 years.^21^ However, registry data on incidence of childhood cancer have only been published for the diagnostic period 1995-2005.^22,23^ Trends in the incidence of childhood cancer over the whole period have not yet been analysed.

This study is the first to assess long-term trends in incidence of childhood cancer in Switzerland. The two major aims are to calculate age-standardised, age-specific, and cumulative incidence for three diagnostic periods—1985-1994, 1995-2004, and 2005-2014—and to describe trends in age-standardised incidence over the entire period from 1985 to 2014. We also assessed indices of quality of registered cases in the SCCR.

## Methods

### Study population and procedure

The Swiss Childhood Cancer Registry (SCCR) was founded by the Swiss Paediatric Oncology Group (SPOG) in 1976 and initially registered patients included in clinical trials.^21^ Beginning in 1981, all patients treated in paediatric cancer centres were systematically included. In a retrospective study covering the period 1990-2004, we discovered, however, that 16% of children with cancer had not been treated in paediatric cancer centres, including infants who were diagnosed only at autopsy in neonatal departments.^24^

Between 2004 and 2007, the SCCR was reorganized according to international recommendations,^25,26^ and the review of diagnoses using pathology reports, quality indices, and retrospective inclusion of missed cases were introduced. Since then, the SCCR improved case completeness by comparing datasets with the population-based cancer registries of each Swiss canton, and by finding missed cases in mortality statistics.^27^ The SCCR includes >95% of all childhood cancer cases diagnosed in Switzerland since 1995.

A clinical research coordinator in each paediatric cancer centre notifies the SCCR of new cases within two months of diagnosis. Children diagnosed with cancer before they reach the age of 16 years and older adolescents diagnosed with typical paediatric tumours are usually treated in paediatric cancer centres in Aarau, Basel, Bellinzona, Bern, Geneva, Lausanne, Lucerne, St. Gallen, and Zurich.

For this study, we included all children with Swiss residency at diagnosis who were registered in the SCCR and diagnosed with cancer according to one of the 12 main diagnostic groups of the ICCC-3 at ages 0-14 years between 1 January 1985 and 31 December 2014.

### Measurements

The SCCR registers patients with leukaemias and lymphomas, malignant and benign CNS tumours, malignant solid tumours, and Langerhans cell histiocytosis who were diagnosed at ages under 21 years in Switzerland. The registry records personal information including name, sex, date of birth, address, nationality; diagnostic information including exact diagnosis and its date, basis of diagnosis, site and grade of primary and subsequent tumour(s), histological type and grade, immunophenotype and genetics, stage at diagnosis, laterality, and site of any metastasis; and treatment information that includes the name and type of treating institution, treatment type, study type, and study protocol name. Diagnoses are classified according to the International Classification of Diseases, tenth revision (ICD-10); topography and morphology of the ICD for Oncology, third revision (ICD-O-3); and the International Classification of Childhood Cancer, third edition (ICCC-3).^25,28,29^ The study was covered by the Ethics Committee of the Canton of Bern approval to the SCCR (KEK-BE: 166/2014).

### Statistical methods

We calculated incidence rates based on population data from the Swiss Federal Statistical Office.^30^ To assess indices of quality, we identified cancers treated in paediatric cancer centres and cancers that had microscopically verified diagnoses that were confirmed from histology of a primary tumour, haematological examination of peripheral blood or bone marrow, or histology of metastasis. We then calculated the average number of cases diagnosed per year and the proportional distribution of ICCC-3 main groups during the three decades 1985-1994, 1995-2004, and 2005-2014. We calculated incidence rates (IR) for ICCC-3 main groups and subgroups per decade expressed per million children,^25^ with direct standardization for age using the 2010 European standard population.^31^ We calculated cumulative IR, defined as the sum of age-specific IR over each year of age from 0 to 14 years and the risk of being diagnosed with cancer before the age of 15 (1 divided by the cumulative IR). We also computed age-specific IRs for the age groups <1, 1-4, 5-9, and 10-14 years. The 95% confidence intervals (95% CI) for the IR were calculated assuming Poisson distribution. We used R version 3.2.2 for data preparation, descriptive statistics, and calculation of IRs. We examined trends in annual age-standardised incidence ratios from 1985 to 2014 for all diagnostic groups combined, separately for boys and girls, and for main diagnostic groups using JoinPoint, Version 4.0.2.2, assessing the magnitude and direction of trends over time and quantifying the average annual percentage change (AAPC).^11,13,16,19,32-35^ JoinPoint fitted straight regression lines through the data, with the natural logarithm of age-standardised incidence rates as dependent variable and calendar year (1985-2014) as independent variable. We used a maximum of two joinpoints to detect a maximum of three different trends, with a minimum of three years between joinpoints. We selected the trend lines that provided the best fit to observed age-standardised incidence rates based on the ratio of the sums of squared errors from the null model and the alternative model. Inference is conducted through Monte Carlo permutation tests as implemented in the software.^33^

The study population included 98 patients (2%) who were registered from death certificate notifications. We had no information on ICCC-3 main group for 12 of these and lacked dates of diagnosis for 75. We classified patients with missing diagnoses as having had other malignant neoplasms and imputed dates of diagnosis with the missforest package for R.^36^ To generate the missing values we used observed data for year of birth, sex, cancer diagnosis, and age at death in the imputation model. We excluded the 98 cases from all analyses using the specific subgroups of ICCC-3 cancer diagnoses (Supplemental Table 1).

## Results

In 2014 Switzerland had a population of 7.825 million inhabitants, of which 1.225 million were 14 years old or younger. From 1985-2014, the SCCR registered 5486 cancer cases in children aged 0-14 years living in Switzerland (Supplemental Figure 1). Of those, 5104 (93%) were microscopically verified (Table 1). The proportion of verified cases of leukaemias has remained at 100% since 1995. Microscopic verification of cancer was highest in 1995-2004 and then decreased again in 2005-2014. This was due to decreases in CNS tumours, retinoblastomas, and renal tumours.

**Table 1.** Numbers of incident cancers and proportions of microscopically verified cancers among all children diagnosed with cancer in Switzerland at age 0-14 years from 1985-2014, by diagnostic period

|  | 1985-1994 |  |  |  | 1995-2004 |  |  |  | 2005-2014 |  |  |  | p-value<sup>b</sup> |
| --- | --- | --- | --- | --- | --- | --- | --- | --- | --- | --- | --- | --- | --- |
|  | All cases | Microscopically verified<sup>a</sup> | % | 95% CI | All cases | Microscopically verified<sup>a</sup> | % | 95% CI | All cases | Microscopically verified<sup>a</sup> | % | 95% CI |  |
| *All cancers (N=5486)* | 1673 | 1539 | 92.0 | 90.7; 93.3 | 1880 | 1769 | 94.1 | 93.0; 95.2 | 1933 | 1796 | 92.9 | 91.8; 94.1 | 0.047 |
| *Age at diagnosis* |  |  |  |  |  |  |  |  |  |  |  |  |  |
| 0-4 years | 804 | 727 | 90.4 | 88.4; 92.5 | 825 | 754 | 91.4 | 89.5; 93.3 | 830 | 755 | 91.0 | 89.0; 92.9 | 0.792 |
| 5-9 years | 444 | 410 | 92.3 | 89.9; 94.8 | 498 | 475 | 95.4 | 93.5; 97.2 | 509 | 472 | 92.7 | 90.5; 95.0 | 0.112 |
| 10-14 years | 425 | 402 | 94.6 | 92.4; 96.7 | 557 | 540 | 96.9 | 95.5; 98.4 | 594 | 569 | 95.8 | 94.2; 97.4 | 0.182 |
| *ICCC-3 Main diagnostic group* |  |  |  |  |  |  |  |  |  |  |  |  |  |
| I. Leukaemias | 569 | 561 | 98.6 | 97.6; 99.6 | 581 | 581 | 100 |  | 647 | 647 | 100 |  | <0.001 |
| II. Lymphomas | 201 | 200 | 99.5 | 98.5; 99.9 | 225 | 225 | 100 |  | 214 | 213 | 99.5 | 98.6; 99.9 | 0.580 |
| III. CNS tumours | 328 | 275 | 83.8 | 79.9; 87.8 | 429 | 376 | 87.6 | 84.5; 90.8 | 448 | 364 | 81.3 | 77.6; 84.9 | 0.033 |
| IV. Neuroblastoma | 134 | 117 | 87.3 | 81.7; 92.9 | 115 | 106 | 92.2 | 87.3; 97.1 | 119 | 112 | 94.1 | 89.9; 98.3 | 0.147 |
| V. Retinoblastoma | 46 | 19 | 41.3 | 27.1; 55.5 | 53 | 17 | 32.1 | 19.5; 44.6 | 42 | 6 | 14.3 | 3.7; 24.9 | 0.020 |
| VI. Renal tumours | 102 | 99 | 97.1 | 93.8; 99.9 | 98 | 97 | 99.0 | 97.0; 99.9 | 98 | 92 | 93.9 | 89.1; 98.6 | 0.135 |
| VII. Hepatic tumours | 18 | 12 | 66.7 | 44.9; 88.4 | 27 | 27 | 100 |  | 16 | 15 | 94 | 81.9; 99.9 | 0.002 |
| VIII. Bone tumours | 73 | 71 | 97 | 93.5; 99.9 | 95 | 95 | 100 |  | 79 | 79 | 100 |  | 0.091 |
| IX. Soft tissue sarcomas | 111 | 110 | 99 | 97.3; 99.9 | 129 | 128 | 99.2 | 97.7; 99.9 | 139 | 139 | 100 |  | 0.555 |
| X. Germ cell tumours | 45 | 42 | 93.3 | 86.0; 99.9 | 53 | 52 | 98.1 | 94.5; 99.9 | 57 | 56 | 98.2 | 94.8; 99.9 | 0.303 |
| XI. Epithelial neoplasms & melanomas | 31 | 29 | 93.5 | 84.9; 99.9 | 65 | 63 | 96.9 | 92.7; 99.9 | 69 | 69 | 100 |  | 0.140 |
| XII. Other malignant neoplasms | 15 | 4 | 26.7 | 4.3; 49.0 | 10 | 2 | 20.0 | 0.0; 44.8 | 5 | 4 | 80 | 44.9; 99.9 | 0.055 |

**Abbreviations:** CI, confidence interval; CNS, central nervous system; ICCC-3, International Classification of Childhood Cancer, third edition.
<sup>a</sup>Includes histology of primary tumours and metastasis, and cytology; <sup>b</sup>p-value derived from Kruskal-Wallis rank sum test for trend, comparing the numbers of microscopically verified cases between the three time periods.

The proportion of cancers treated in paediatric cancer centres increased from 84% during 1985-1994 to 98% for 2005-2014 (Supplemental Table 2). The increase was largest for CNS tumours and epithelial neoplasms and melanomas; the proportion of CNS tumours treated in paediatric cancer centres increased from 71% in 1985-1994 to 98% in 2005-2014.

During the decade 1985-1994, 1673 cancer cases were diagnosed at age 0-14 years; 1880 cases were diagnosed in 1995-2004, and 1933 cases in 2005-2014 (Table 1). In all three decades, the most common cancers were leukaemias (31-34%), CNS tumours (19-23%), and lymphomas (11-12%) (Table 2). The overall annual age-standardised incidence rate per million children rose from 143 for 1985-1994 to 162 during 2005-2014 (Table 2, Figure 1). Age-standardised incidence in all three decades was highest for leukaemias, CNS tumours, and lymphomas, and the overall cumulative incidence before the age of 14 years rose from 2135 to 2423 per million children from 1985-1994 to 2005-2014.

**Table 2.** Incidence of childhood cancer in Switzerland at age 0-14 years from 1985-2014, by diagnostic group and period

| 1985-1994 |  |  |  |  |  |
| --- | --- | --- | --- | --- | --- |
|  | Average number<br>of cases/year (%) | ASR per million<br>population/year<br>(95% CI)<sup>a</sup> | CR per million<br>population<br>(95% CI)<sup>b</sup> | 1:N<br>children<sup>c</sup> | Male/<br>Female<sup>d</sup> |
| *All cancers* | 166.6 (100) | 143.1 (136.2; 150.0) | 2135.3 (2108.8; 2161.8) | 468 | 1.24 |
| *ICCC-3 Main diagnostic group* |  |  |  |  |  |
| I. Leukaemias | 56.9 (34.2) | 48.8 (44.8; 52.8) | 730.3 (714.8; 745.8) | 1369 | 1.29 |
| II. Lymphomas | 20.1 (12.1) | 17.3 (14.9; 19.7) | 259.2 (249.9; 268.4) | 3858 | 2.14 |
| III. CNS tumours | 32.4 (19.4) | 27.8 (24.8; 30.8) | 417.1 (405.4; 428.8) | 2398 | 1.17 |
| IV. Neuroblastoma | 13.4 (8.0) | 11.5 (9.6; 13.5) | 168.9 (161.4; 176.3) | 5921 | 1.27 |
| V. Retinoblastoma | 4.6 (2.8) | 4.0 (2.8; 5.1) | 58.0 (53.6; 62.4) | 17241 | 0.84 |
| VI. Renal tumours | 10.2 (6.1) | 8.8 (7.1; 10.5) | 129.8 (123.3; 136.4) | 7704 | 0.76 |
| VII. Hepatic tumours | 1.8 (1.1) | 1.6 (0.8; 2.3) | 22.9 (20.1; 25.6) | 43668 | 1.25 |
| VIII. Bone tumours | 7.3 (4.4) | 6.3 (4.8; 7.7) | 94.3 (88.7; 99.9) | 10604 | 1.03 |
| IX. Soft tissue sarcomas | 10.9 (6.5) | 9.4 (7.6; 11.1) | 139.6 (132.8; 146.3) | 7163 | 1.06 |
| X. Germ cell tumours | 4.4 (2.6) | 3.8 (2.7; 4.9) | 56.4 (52.1; 60.7) | 17730 | 1.10 |
| XI. Epithelial neoplasms & melanomas | 3.1 (1.9) | 2.7 (1.7; 3.6) | 39.9 (36.3; 43.5) | 25063 | 1.07 |
| XII. Other malignant neoplasms | 1.5 (0.9) | 1.3 (0.6; 1.9) | 19.1 (16.6; 21.6) | 52356 | 0.88 |
| 1995-2004 |  |  |  |  |  |
|  | Average number<br>of cases/year (%) | ASR per million<br>population/year<br>(95% CI)<sup>a</sup> | CR per million<br>population<br>(95% CI)<sup>b</sup> | 1:N<br>children<sup>c</sup> | Male/<br>Female<sup>d</sup> |
| *All cancers* | 187.5 (100) | 153.8 (146.8; 160.8) | 2298.9 (2272.2; 2325.6) | 435 | 1.27 |
| *ICCC-3 Main diagnostic group* |  |  |  |  |  |
| I. Leukaemias | 57.8 (30.8) | 47.4 (43.6; 51.3) | 710.3 (695.4; 725.1) | 1408 | 1.40 |
| II. Lymphomas | 22.5 (12.0) | 18.1 (15.7; 20.4) | 270.6 (261.4; 279.7) | 3695 | 1.81 |
| III. CNS tumours | 42.9 (22.9) | 34.8 (31.5; 38.1) | 523.4 (510.6; 536.1) | 1911 | 1.25 |
| IV. Neuroblastoma | 11.3 (6.0) | 9.9 (8.0; 11.7) | 144.6 (137.9; 151.3) | 6916 | 0.92 |
| V. Retinoblastoma | 5.3 (2.8) | 4.7 (3.4; 5.9) | 68.4 (63.7; 73.0) | 14620 | 0.96 |
| VI. Renal tumours | 9.8 (5.2) | 8.2 (6.6; 9.9) | 122.5 (116.3; 128.6) | 8163 | 1.00 |
| VII. Hepatic tumours | 2.7 (1.4) | 2.3 (1.4; 3.2) | 33.9 (30.6; 37.1) | 29499 | 3.50 |
| VIII. Bone tumours | 9.5 (5.1) | 7.5 (6.0; 9.0) | 112.8 (106.9; 118.7) | 8865 | 0.94 |
| IX. Soft tissue sarcomas | 12.9 (6.9) | 10.6 (8.7; 12.4) | 157.7 (150.7; 164.7) | 6341 | 1.48 |
| X. Germ cell tumours | 5.3 (2.8) | 4.4 (3.2; 5.5) | 65.1 (60.6; 69.5) | 15361 | 0.77 |
| XI. Epithelial neoplasms & melanomas | 6.5 (3.5) | 5.2 (3.9; 6.4) | 77.5 (72.5; 82.4) | 12903 | 0.81 |
| XII. Other malignant neoplasms | 1.0 (0.5) | 0.8 (0.3; 1.4) | 12.3 (10.4; 14.3) | 81301 | 1.00 |
| 2005-2014 |  |  |  |  |  |
|  | Average number<br>of cases/year (%) | ASR per million<br>population/year<br>(95% CI)<sup>a</sup> | CR per million<br>population<br>(95% CI)<sup>b</sup> | 1:N<br>children<sup>c</sup> | Male/<br>Female<sup>d</sup> |
| *All cancers* | 192.4 (100) | 162.3 (155.0; 169.5) | 2423.4 (2395.5; 2451.2) | 413 | 1.28 |
| *ICCC-3 Main diagnostic group* |  |  |  |  |  |
| I. Leukaemias | 64.3 (33.4) | 54.5 (50.3; 58.8) | 816.0 (799.8; 832.2) | 1225 | 1.57 |
| II. Lymphomas | 21.3 (11.1) | 17.5 (15.2; 19.9) | 262.4 (253.2; 271.5) | 3811 | 1.77 |
| III. CNS tumours | 44.5 (23.1) | 37.3 (33.9; 40.8) | 560.7 (547.3; 574.1) | 1783 | 1.17 |
| IV. Neuroblastoma | 11.9 (6.2) | 10.5 (8.6; 12.3) | 153.0 (146.0; 160.0) | 6536 | 1.13 |
| V. Retinoblastoma | 4.2 (2.2) | 3.7 (2.6; 4.8) | 54.1 (49.9; 58.3) | 18484 | 0.83 |
| VI. Renal tumours | 9.7 (5.0) | 8.4 (6.7; 10.0) | 124.4 (118.1; 130.7) | 8039 | 0.80 |
| VII. Hepatic tumours | 1.6 (0.8) | 1.4 (0.7; 2.1) | 20.4 (17.9; 23.0) | 49020 | 1.67 |
| VIII. Bone tumours | 7.9 (4.1) | 6.5 (5.0; 7.9) | 97.1 (91.5; 102.7) | 10299 | 0.88 |
| IX. Soft tissue sarcomas | 13.9 (7.2) | 11.6 (9.7; 13.6) | 173.9 (166.5; 181.4) | 5750 | 1.36 |
| X. Germ cell tumours | 5.7 (3.0) | 4.8 (3.6; 6.1) | 71.0 (66.2; 75.8) | 14085 | 1.19 |
| XI. Epithelial neoplasms & melanomas | 6.9 (3.6) | 5.6 (4.3; 7.0) | 84.0 (78.8; 89.2) | 11905 | 0.60 |
| XII. Other malignant neoplasms | 0.5 (0.3) | 0.4 (0.1; 0.8) | 6.3 (4.9; 7.7) | 158730 | 1.50 |

**Abbreviations:** ASR, age-standardised incidence rate; CI, confidence interval; CNS, central nervous system; CR, cumulative incidence rate; ICCC-3, International Classification of Childhood Cancer, third edition.
<sup>a</sup>Standardised according to the 2010 European standard population; <sup>b</sup>Cumulative incidence up to the age of 14 years; <sup>c</sup>Number of children affected up to the age of 14 years in Switzerland; <sup>d</sup>male:female ratio

**Figure 1.**
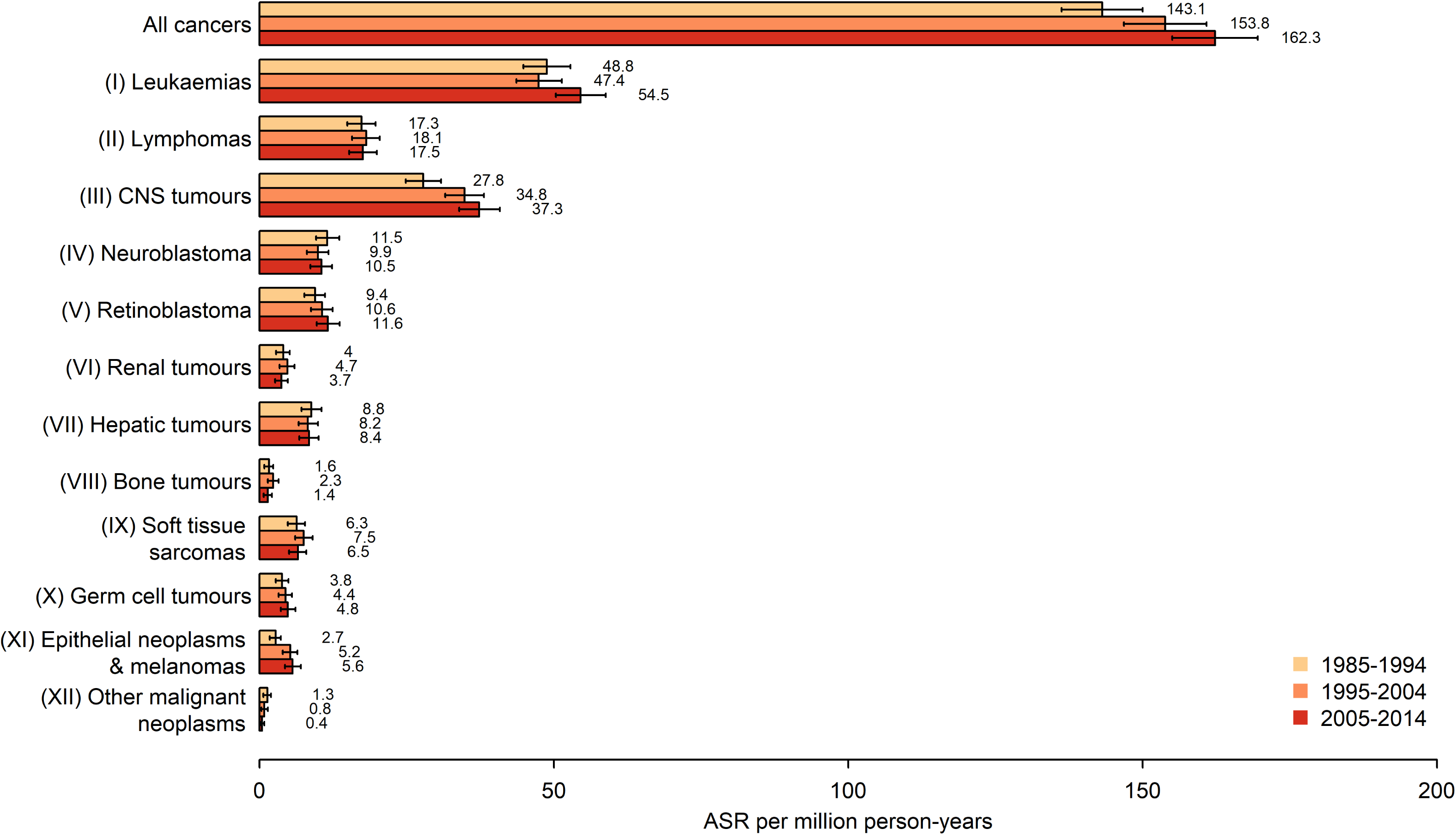
Comparison of incidence of childhood cancer between 1985-1994, 1995-2004 and 2005-2014, by cancer diagnosis. Abbreviations: ASR, Age-standardized incidence rate. Cancer diagnoses were classified according to the International Classification of Childhood Cancer, third edition (ICCC-3). Incidence rates were age-standardized according to the 2010 European standard population. Error bars show 95% confidence intervals.

Childhood cancer was more common in boys than in girls, with a male to female ratio of 1.28 during 2005-2014 (Table 2). The male to female ratio was largest for Burkitt’s lymphoma, 6.71, and smallest for thyroid carcinomas, 0.27 (Supplemental Table 1). Table 3 lists the age-specific incidences of childhood cancer over the period 2005-2014. Leukaemias and CNS tumours were among the most common cancers throughout all age group, neuroblastoma until 4 years, renal tumours from 1-4 years, and lymphoma from 5-14 years.

**Table 3.** Age-specific incidence rate per million population in Switzerland and average cases per year from 2005-2014, by age at diagnosis and diagnostic group

|  | Age-specific incidence rate (95% CI) |  |  |  | Average cases per year per million population |  |  |  |
| --- | --- | --- | --- | --- | --- | --- | --- | --- |
|  | <1 year | 1-4 years | 5-9 years | 10-14 years | <1 year | 1-4 years | 5-9 years | 10-14 years |
| *All cancers* | 228.3 (195.9; 264.6) | 209.5 (193.7; 226.3) | 129.7 (118.6; 141.5) | 142.0 (130.8; 154.0) | 11.8 | 43.4 | 33.7 | 39.4 |
| *ICCC-3 Main diagnostic group* |  |  |  |  |  |  |  |  |
| I. Leukaemias | 36.1 (24.0; 52.2) | 99.1 (88.4; 110.8) | 43.9 (37.6; 51.0) | 32.7 (27.4; 38.7) | 1.9 | 20.5 | 11.4 | 9.1 |
| II. Lymphomas | 3.9 (0.8; 11.3) | 9.7 (6.5; 13.8) | 14.9 (11.3; 19.3) | 29.3 (24.4; 35.0) | 0.2 | 2.0 | 3.9 | 8.1 |
| III. CNS tumours | 36.1 (24.0; 52.2) | 38.3 (31.7; 45.8) | 40.6 (34.5; 47.4) | 33.6 (28.3; 39.7) | 1.9 | 7.9 | 10.5 | 9.3 |
| IV. Neuroblastoma | 65.8 (49.0; 86.5) | 18.7 (14.2; 24.1) | 1.5 (0.6; 3.4) | 1.0 (0.3; 2.5) | 3.4 | 3.9 | 0.4 | 0.3 |
| V. Retinoblastoma | 25.8 (15.8; 39.8) | 6.4 (3.9; 9.9) | 0.5 (0.1; 1.9) | 0.0 (0.0; 0.7) | 1.3 | 1.3 | 0.1 | 0.0 |
| VI. Renal tumours | 18.1 (9.9; 30.3) | 16.7 (12.5; 21.9) | 6.4 (4.2; 9.5) | 1.4 (0.5; 3.1) | 0.9 | 3.5 | 1.7 | 0.4 |
| VII. Hepatic tumours | 7.7 (2.8; 16.8) | 2.3 (0.9; 4.6) | 0.5 (0.1; 1.9) | 0.2 (0.0; 1.3) | 0.4 | 0.5 | 0.1 | 0.1 |
| VIII. Bone tumours | 0.0 (0.0; 3.9) | 1.6 (0.5; 3.8) | 5.6 (3.5; 8.6) | 12.5 (9.3; 16.4) | 0.0 | 0.3 | 1.5 | 3.5 |
| IX. Soft tissue sarcomas | 11.6 (5.3; 22.0) | 12.6 (8.9; 17.2) | 9.2 (6.5; 12.8) | 13.2 (10.0; 17.2) | 0.6 | 2.6 | 2.4 | 3.7 |
| X. Germ cell tumours | 19.4 (10.8; 31.9) | 3.5 (1.8; 6.3) | 1.8 (0.7; 3.7) | 5.8 (3.7; 8.6) | 1.0 | 0.7 | 0.5 | 1.6 |
| XI. Epithelial neoplasms & melanomas | 1.3 (0.0; 7.2) | 0.3 (0.0; 1.8) | 4.4 (2.5; 7.0) | 12.0 (8.9; 15.8) | 0.1 | 0.1 | 1.1 | 3.3 |
| XII. Other malignant neoplasms | 2.6 (0.3; 9.3) | 0.3 (0.0; 1.8) | 0.3 (0.0; 1.4) | 0.2 (0.0; 1.3) | 0.1 | 0.1 | 0.1 | 0.1 |

**Abbreviations:** CI, confidence interval; CNS, central nervous system; ICCC-3, International Classification of Childhood Cancer, third edition.

Overall, age-standardised childhood cancer incidence increased between 1985-2014 by 0.7% per year (95% CI 0.5%-1.0%), though trends differed between age groups, type of cancer, and sex (Table 4, Figure 2). This increase was highest for children diagnosed between the ages of 10 and 14 years. The annual increase of 0.8% (95% CI 0.2%-1.4%) from 1985-2014 for the incidence of leukaemias was mainly driven by a 1.2% (95% CI 0.4%-2.0%) increase among boys (Table 4). The incidence of CNS tumours increased annually by 3.0% (95% CI 1.3%-4.6%) from 1985-2002, but not thereafter (Table 4). Over the entire period, to 2014, the annual increase was 1.4% (95% CI 0.5%-2.4%) for boys and 1.7% (95% CI 0.7%-2.7%) for girls. Incidence of epithelial neoplasms and melanomas increased by 3.8% (95% CI 1.7%-6.0%) from 1985-2014, but because they account for only 3% of all cancer cases (165 of 5486 total cases) this had a very small effect on the overall trend (Tables 1 and 4).

**Figure 2.**
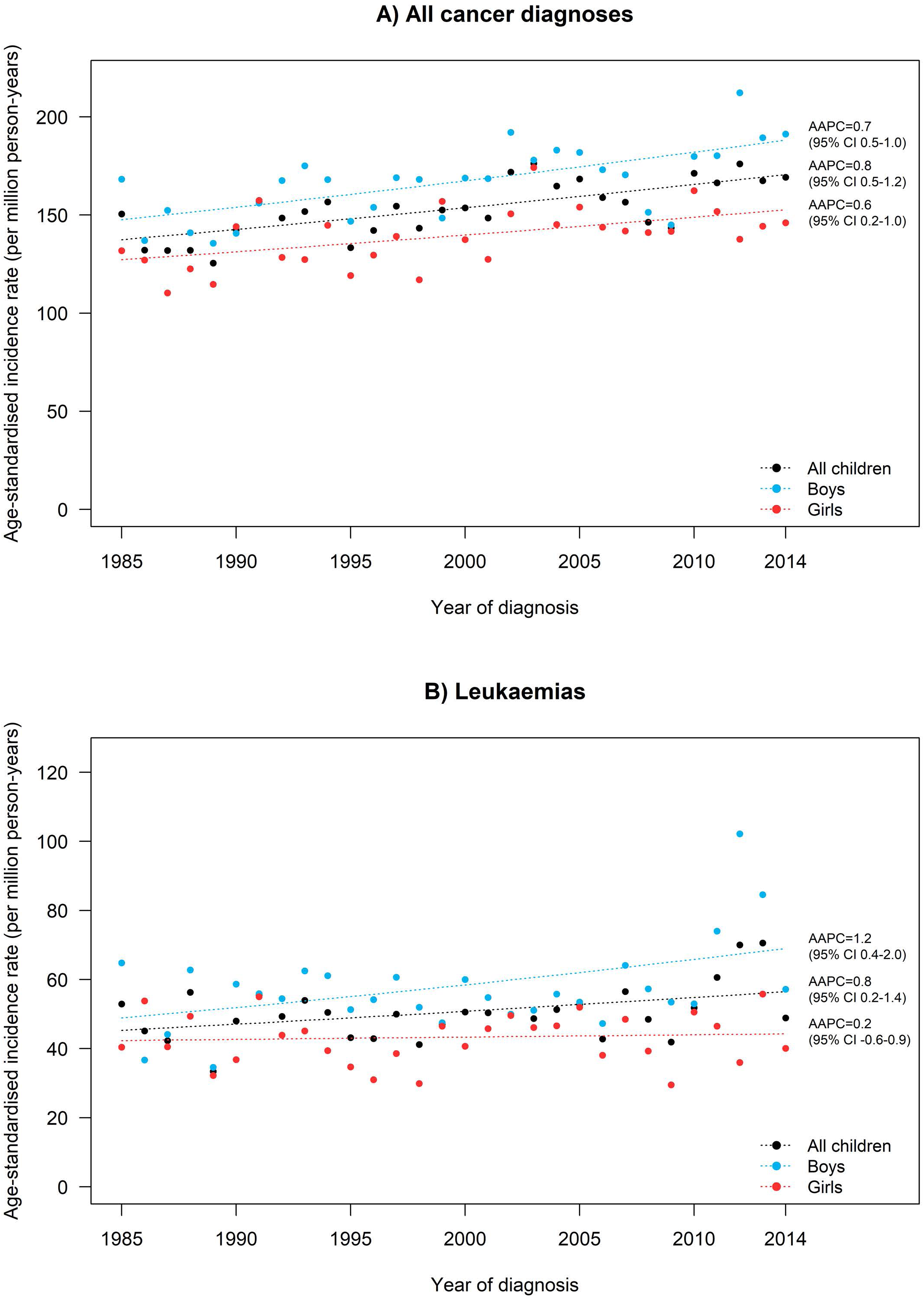
Trends in incidence rates (per million person-years) in Switzerland between 1985 and 2014, standardized according to the 2010 European standard population, for A) all childhood cancer diagnoses combined and B) leukaemias. Trends were modelled using Joinpoint regression. Abbreviations: AAPC, average annual percentage change; CI, confidence interval.

**Table 4.** Average annual percentage change (AAPC) in incidence rates of childhood cancer by diagnostic group, age and sex in Switzerland 1985-2014: Results from JoinPoint regression.

| Diagnostic group | Trend 1 |  | Trend 2 |  |
| --- | --- | --- | --- | --- |
|  | Period | AAPC (95% CI) | Period | AAPC (95% CI) |
| *All cancers* | 1985-2014 | **0.7 (0.5; 1.0)** |  |  |
| 0-4 years | 1985-2014 | 0.3 (-0.1; 0.7) |  |  |
| 5-9 years | 1985-2014 | **0.8 (0.2; 1.3)** |  |  |
| 10-14 years | 1985-2014 | **1.4 (0.9; 2.0)** |  |  |
| I. Leukaemias | 1985-2009 | **0.8 (0.2; 1.4)** |  |  |
| II. Lymphomas | 1985-2014 | 0.0 (-0.9; 1.0) |  |  |
| III. CNS tumours | 1985-2002 | **3.0 (1.3; 4.6)** | 2002-2014 | -0.2 (-2.4; 2.1) |
| IV. Neuroblastoma | 1985-2014 | -0.5 (-1.7; 0.7) |  |  |
| V. Retinoblastoma | 1985-2014 | **-0.9 (-3.0; 1.2)** |  |  |
| VI. Renal tumours | 1985-2014 | -0.3 (-1.5; 0.9) |  |  |
| VII. Hepatic tumours | 1985-2014 | n.a. |  |  |
| VIII. Bone tumours | 1985-2014 | 0.2 (-0.9; 1.4) |  |  |
| IX. Soft tissue sarcomas | 1985-2014 | 1.0 (-0.4; 2.5) |  |  |
| X. Germ cell tumours | 1985-2014 | 1.6 (-0.2; 3.4) |  |  |
| XI. Epithelial neoplasms & melanomas | 1985-2014 | **3.8 (1.7; 6.0)** |  |  |
| XII. Other malignant neoplasms | 1985-2014 | n.a. |  |  |
| *Boys* |  |  |  |  |
| All cases | 1985-2014 | **0.8 (0.5; 1.2)** |  |  |
| I. Leukaemias | 1985-2010 | **1.2 (0.4; 2.0)** |  |  |
| II. Lymphomas | 1985-2014 | 0.0 (-1.1; 1.0) |  |  |
| III. CNS tumours | 1985-2014 | **1.4 (0.5; 2.4)** |  |  |
| IV. Neuroblastoma | 1985-2014 | -1.1 (-2.3; 0.1) |  |  |
| V. Retinoblastoma | 1985-2014 | n.a. |  |  |
| VI. Renal tumours | 1985-2014 | 0.1 (-1.9; 2.1) |  |  |
| VII. Hepatic tumours | 1985-2014 | n.a. |  |  |
| VIII. Bone tumours | 1985-2014 | n.a. |  |  |
| IX. Soft tissue sarcomas | 1985-2014 | 0.7 (-0.8; 2.2) |  |  |
| X. Germ cell tumours | 1985-2014 | n.a. |  |  |
| XI. Epithelial neoplasms & melanomas | 1985-2014 | n.a. |  |  |
| XII. Other malignant neoplasms | 1985-2014 | n.a. |  |  |
| *Girls* |  |  |  |  |
| All cases | 1985-2014 | **0.6 (0.2; 1.0)** |  |  |
| I. Leukaemias | 1985-2014 | 0.2 (-0.6; 0.9) |  |  |
| II. Lymphomas | 1985-2014 | 0.1 (-1.8; 2.0) |  |  |
| III. CNS tumours | 1985-2014 | **1.7 (0.7; 2.7)** |  |  |
| IV. Neuroblastoma | 1985-2014 | 0.6 (-1.3; 2.6) |  |  |
| V. Retinoblastoma | 1985-2014 | n.a. |  |  |
| VI. Renal tumours | 1985-2014 | -0.7 (-2.4; 1.1) |  |  |
| VII. Hepatic tumours | 1985-2014 | n.a. |  |  |
| VIII. Bone tumours | 1985-2014 | 0.7 (-0.9; 2.5) |  |  |
| IX. Soft tissue sarcomas | 1985-2014 | n.a. |  |  |
| X. Germ cell tumours | 1985-2014 | n.a. |  |  |
| XI. Epithelial neoplasms & melanomas | 1985-2014 | n.a. |  |  |
| XII. Other malignant neoplasms | 1985-2014 | n.a. |  |  |

**Abbreviations:** AAPC, average annual percentage change; CI, confidence interval; CNS, central nervous system; n.a., not applicable.

## Discussion

Overall childhood cancer incidence rates significantly increased between 1985-2014 in Switzerland, mainly driven by leukaemias and CNS tumours. The trend for CNS tumours plateaued in the 2000s, but continued for leukaemias and epithelial neoplasms and melanomas. The increase in overall incidence rates was age-dependent, with no increase in preschool children up to age 4, a slight increase in children 5-9 years old, and a stronger increase in young adolescents 10-14 years old.

### International comparison

The proportion of microscopically verified cases in the SCCR is high (>93%), suggesting excellent validity of cancer diagnoses. This is comparable to validity ranging from 92-98% reported in other European countries,^17,37,12,1,38,11^ and in Japan (90%),^39^ the US (95%),^32^ and Australia (97%).^13^ Lower proportions of microscopically verified cases in certain tumours do not necessarily mean lower data quality, though. For retinoblastoma, current treatment rarely involves surgical removal of the eye, which has led to a decreasing proportion of microscopically verified cases from 41% for 1985-1994 to 14% during 2005-2014. Patients with low-grade gliomas and certain high-grade tumours are increasingly diagnosed with imaging techniques when surgical intervention puts them at high risk for functional loss; many of these patients do not undergo any surgery unless it becomes inevitable. The decreasing proportion of microscopically verified CNS tumours, from 92% during 1985-1994 to 82% during 2005-2014, reflects this.

Supplemental Table 3 compares the relative distribution of diagnostic groups and the overall age-standardised incidence of childhood cancer in Switzerland to the Piedmont region in Italy,^11^ Spain,^12^ Sweden,^9^ France,^37^ Germany,^38^ Austria,^17^ Western Europe combined,^1^ USA,^40^ Canada,^41^ Australia,^13^ and Korea.^14^ The relative distribution of diagnostic groups was similar to these countries, but age-standardized leukaemia incidence was among the highest, with higher values observed in only Sweden, Germany, and Australia. The incidence of neuroblastoma was among the lowest, excepting only Sweden and Australia.

Overall cancer incidence rates in Switzerland continuously rose over the whole study period, from 1985-2014. Overall incidence rates increased in all world regions except sub-Saharan Africa from the 1980s to 2001-2010,^1^ but have remained stable in Austria,^17^ Ireland,^18^ the USA,^19^ Canada,^16^ and Australia^13^ in more recent time periods (Supplemental Table 4). The increased incidence of childhood cancer observed in our study is mainly attributable to leukaemia and CNS tumours. Incidence of leukaemia increased steadily until 2014 without evidence of a plateau. Most countries also observed an increase,^2,7-9,11-14,18,20^ while studies from the USA and Austria reported stable leukaemia incidence (Supplemental Table 4).^17,19,32^ Leukaemia incidence in Canada first increased but levelled off after 1999.^16^ We observed that incidence of CNS tumours increased until 2002 and plateaued thereafter, similar to Australia.^13^ Many studies found increases over the entire study period,^8,11,12,17,20^ others found no increase,^14,16,19,32,35,42,43^ or even a decrease.^35^ We also found that incidence of epithelial neoplasms and melanomas increased over the entire study period, similar to the USA,^10,32^ Australia,^13^ and Korea.^14^

### Possible reasons for increasing trend in incidence

The increased incidence of childhood cancer may be attributable to several factors. First, improved case registration would increase the observed incidence. Patients with CNS tumours, with epithelial neoplasms and melanomas, and aged 10-14 years at time of cancer diagnosis increasingly have been treated in paediatric oncology centres. These paediatric oncology centres collaborate closely with the SCCR and actively report every newly diagnosed child.^24^ Since 1985, SCCR coverage has improved from 85% to 95%.^27^ The SCCR has retrospectively registered missed cases, but we cannot exclude that some of the children with CNS tumours or children treated in earlier decades in adult facilities escaped registration, and thus that earlier incidence of childhood cancers is underestimated. Advances in medical diagnostics also may have increased apparent incidence rates. Imaging techniques have enhanced the capacity to diagnose otherwise undetected low-grade CNS tumours since the mid-1980s.^2,44^ Finally, medical and environmental risk factors could have increased the incidence of some cancers, including leukaemias in particular, which recently have increased in nearly all countries. Leukaemia and CNS tumours may be associated with exposure to low-level ionizing radiation,^45^ which has likely increased in recent decades due to the increasing use of imaging techniques that rely upon ionizing radiation, especially computed tomography.^46^ This might explain the age-dependent pattern of increase. Other potential risk factors include genetic causes, higher birth weight and parental age, increased infections, exposure to pesticides and traffic-related air pollution, as well as parental exposure to benzene.^47-49^ However, the results of this study do not allow us to draw conclusions about specific risk factors that may have contributed to the overall increasing incidence of cancer in general, and specifically of leukaemia and CNS tumours.

### Strengths and limitations of the study

The nationwide, population-based coverage of the SCCR with a high completeness of registration is a real strength of this study.^27^ Data quality was high with more than 90% of the cases having been microscopically verified. We could include recent diagnostic years (until 2014) due to the fast reporting and quality control procedures in the SCCR.^21^ For the most recent years, the SCCR is in the process of exchanging data with the cantonal cancer registries. Therefore, we may have underestimated increases in incidence in the most recent period, although the SCCR missed very few cases in previous linkages. The sample size was limited by the comparatively small population of Switzerland, which led to wide confidence intervals for some diagnostic groups. Results should thus be interpreted cautiously because trends may reflect random fluctuations in incidence.

## Conclusions

Changes in registration procedures and advances in medical diagnostics may explain the increase observed in the incidence in CNS tumours. For leukaemia, rising incidence may be real and due to changes in parental lifestyle, infections, pesticide exposure, air pollution, birth weight, or other risk factors. Future aetiological research should examine these long-term trends together with changes in medical and environmental risk factors.

## ACKNOWLEDGEMENTS

We thank the clinical research coordinators of the Swiss Paediatric Oncology Group: Dr. sc. nat. Claudia Althaus, Nadine Amport, Pamela Balestra, Nadine Beusch, Sarah Blanc, Susann Drerup, Janine Garibay, Franziska Hochreutener, Monika Imbach, Friedgard Julmy, Rachel Simone Kusche, Eléna Lemmel, Heike Markiewicz, Dr. med. Veneranda Mattiello, Rodolfo Lo Piccolo, Annette Reinberg, Astrid Schiltknecht, Renate Siegenthaler, Verena Stahel. We also thank the team of the Swiss Childhood Cancer Registry: Meltem Altun, Erika Berclaz-Brantschen, Katharina Flandera, Elisabeth Kiraly. We thank Christopher Ritter for his editorial suggestions and Ben Spycher for statistical advice.

This study was supported by the Swiss National Science Foundation (grant PDFMP3_141775), the Swiss Bridge Foundation (www.swissbridge.ch), the Swiss Cancer League (KLS-3412-02-2014, KLS-3886-02-2016), the Swiss Cancer Research foundation (KFS-4157-02-2017), the Bernese Cancer League and Kinderkrebs Schweiz. The work of the Swiss Childhood Cancer Registry is supported by the Swiss Paediatric Oncology Group (www.spog.ch), Schweizerische Konferenz der kantonalen Gesundheitsdirektorinnen und –direktoren (www.gdk-cds.ch), Swiss Cancer Research (www.krebsforschung.ch), Kinderkrebshilfe Schweiz (www.kinderkrebshilfe.ch), Bundesamt für Gesundheit, NICER and Celgene.

## REFERENCES

1. Steliarova-Foucher E, Colombet M, Ries LAG, Moreno F, Dolya A, Bray F, Hesseling P, Shin HY, Stiller CA, contributors I-. International incidence of childhood cancer, 2001-10: a population-based registry study. Lancet Oncol 2017; 10.1016/S1470-2045(17)30186-9.

2. Kroll ME, Carpenter LM, Murphy MF, Stiller CA. Effects of changes in diagnosis and registration on time trends in recorded childhood cancer incidence in Great Britain. Br J Cancer 2012; 107: 1159–62.

3. Adamson P, Law G, Roman E. Assessment of trends in childhood cancer incidence. Lancet 2005; 365: 753.

4. McNally RJ, Kelsey AM, Cairns DP, Taylor GM, Eden OB, Birch JM. Temporal increases in the incidence of childhood solid tumors seen in Northwest England (1954-1998) are likely to be real. Cancer 2001; 92: 1967–76.

5. McNally RJ, Cairns DP, Eden OB, Kelsey AM, Taylor GM, Birch JM. Examination of temporal trends in the incidence of childhood leukaemias and lymphomas provides aetiological clues. Leukemia 2001; 15: 1612–8.

6. Steliarova-Foucher E, Stiller C, Kaatsch P, Berrino F, Coebergh JW, Lacour B, Parkin M. Geographical patterns and time trends of cancer incidence and survival among children and adolescents in Europe since the 1970s (the ACCISproject): an epidemiological study. Lancet 2004; 364: 2097–105.

7. Kroll ME, Draper GJ, Stiller CA, Murphy MF. Childhood leukemia incidence in Britain, 1974-2000: time trends and possible relation to influenza epidemics. J Natl Cancer Inst 2006; 98: 417–20.

8. Spix C, Eletr D, Blettner M, Kaatsch P. Temporal trends in the incidence rate of childhood cancer in Germany 1987-2004. Int J Cancer 2008; 122: 1859–67.

9. Dreifaldt AC, Carlberg M, Hardell L. Increasing incidence rates of childhood malignant diseases in Sweden during the period 1960-1998. Eur J Cancer 2004; 40: 1351–60.

10. Ward EM, Thun MJ, Hannan LM, Jemal A. Interpreting cancer trends. Ann N Y Acad Sci 2006; 1076: 29–53.

11. Isaevska E, Manasievska M, Alessi D, Mosso ML, Magnani C, Sacerdote C, Pastore G, Fagioli F, Merletti F, Maule M. Cancer incidence rates and trends among children and adolescents in Piedmont, 1967-2011. PLoS One 2017; 12: e0181805.

12. Peris-Bonet R, Salmeron D, Martinez-Beneito MA, Galceran J, Marcos-Gragera R, Felipe S, Gonzalez V, Sanchez de Toledo Codina J. Childhood cancer incidence and survival in Spain. Ann Oncol 2010; 21 Suppl 3: iii103–10.

13. Baade PD, Youlden DR, Valery PC, Hassall T, Ward L, Green AC, Aitken JF. Trends in incidence of childhood cancer in Australia, 1983-2006. Br J Cancer 2010; 102: 620–6.

14. Park HJ, Moon EK, Yoon JY, Oh CM, Jung KW, Park BK, Shin HY, Won YJ. Incidence and Survival of Childhood Cancer in Korea. Cancer Res Treat 2016; 10.4143/crt.2015.290.

15. Zheng R, Peng X, Zeng H, Zhang S, Chen T, Wang H, Chen W. Incidence, mortality and survival of childhood cancer in China during 2000-2010 period: A population-based study. Cancer Lett 2015; 363: 176–80.

16. Mitra D, Shaw AK, Hutchings K. Trends in incidence of childhood cancer in Canada, 1992-2006. Chronic Dis Inj Can 2012; 32: 131–9.

17. Karim-Kos HE, Hackl M, Mann G, Urban C, Woehrer A, Slavc I, Ladenstein R. Trends in incidence, survival and mortality of childhood and adolescent cancer in Austria, 1994-2011. Cancer Epidemiol 2016; 42: 72–81.

18. National Cancer Registry Ireland. Cancer Trends; 2014.

19. Siegel DA, King J, Tai E, Buchanan N, Ajani UA, Li J. Cancer incidence rates and trends among children and adolescents in the United States, 2001-2009. Pediatrics 2014; 134: e945–55.

20. Pesola F, Ferlay J, Sasieni P. Cancer incidence in English children, adolescents and young people: past trends and projections to 2030. Br J Cancer 2017; 117: 1865–73.

21. Pfeiffer V, Redmond S, Kuonen R, Sommer G, Spycher B, Schindler M, Singh P, Michel G, Kuehni C. Swiss Childhood Cancer Registry - Annual Report 2014/2015. Institute of Social and Preventive Medicine, University of Bern, Switzerland; 2015.

22. Michel G, von der Weid NX, Zwahlen M, Adam M, Rebholz CE, Kuehni CE, Swiss Childhood Cancer Registry, Swiss Paediatric Oncology Group. The Swiss Childhood Cancer Registry: rationale, organisation and results for the years 2001-2005. Swiss Med Wkly 2007; 137: 502–9.

23. Michel G, von der Weid NX, Zwahlen M, Redmond S, Strippoli MP, Kuehni CE, Swiss Paediatric Oncology Group. Incidence of childhood cancer in Switzerland: the Swiss Childhood Cancer Registry. Pediatr Blood Cancer 2008; 50: 46–51.

24. Adam M, von der Weid N, Michel G, Zwahlen M, Lutz JM, Probst-Hensch N, Niggli F, Kuehni C, Swiss Pediatric Oncology Group, Swiss Association of Cancer Registries. Access to specialized pediatric cancer care in Switzerland. Pediatr Blood Cancer 2010; 54: 721–7.

25. Steliarova-Foucher E, Stiller C, Lacour B, Kaatsch P. International Classification of Childhood Cancer, third edition. Cancer 2005; 103: 1457–67.

26. Jensen OM, Parkin DM, MacLennan R, Muir CS, Skeet RG. Cancer Registration: Principles and Methods. IARC Sci Publ 1991.

27. Schindler M, Mitter V, Bergstraesser E, Gumy-Pause F, Michel G, Kuehni CE. Death certificate notifications in the Swiss Childhood Cancer Registry: assessing completeness and registration procedures. Swiss Med Wkly 2015; 145: w14225.

28. Fritz A, Percy C, Jack A, Shanmugaratnam K, Sobin L, Parkin D, Whelan S. International Classification of Diseases for Oncology, third edition. Geneva, World Health Organisation; 2000.

29. World Health Organisation. International Statistical Classification of Diseases and Related Health Problems. - 10th Revision. Geneva; 1992.

30. Swiss Federal Statistical Office. Population and Households Statistics (STATPOP). 2016. http://www.bfs.admin.ch/bfs/portal/de/index/news/02/03/01/01.html.

31. European Union. Revision of the European Standard Population: Report of Eurostat’s task force. Luxembourg; 2013.

32. Linabery AM, Ross JA. Trends in childhood cancer incidence in the U.S. (1992-2004). Cancer 2008; 112: 416–32.

33. National Cancer Institute. Joinpoint Trend Analysis Software. 2016. http://surveillance.cancer.gov/joinpoint/.

34. Barrington-Trimis JL, Cockburn M, Metayer C, Gauderman WJ, Wiemels J, McKean-Cowdin R. Trends in childhood leukemia incidence over two decades from 1992 to 2013. Int J Cancer 2017; 140: 1000–8.

35. Papathoma P, Thomopoulos TP, Karalexi MA, Ryzhov A, Zborovskaya A, Dimitrova N, Zivkovic S, Eser S, Antunes L, Sekerija M, Zagar T, Bastos J, Demetriou A, Cozma R, Coza D, Bouka E, Dessypris N, Kantzanou M, Kanavidis P, Dana H, Hatzipantelis E, Moschovi M, Polychronopoulou S, Pourtsidis A, Stiakaki E, Papakonstantinou E, Oikonomou K, Sgouros S, Vakis A, Zountsas B, Bourgioti C, Kelekis N, Prassopoulos P, Choreftaki T, Papadopoulos S, Stefanaki K, Strantzia K, Cardis E, Steliarova-Foucher E, Petridou ET. Childhood central nervous system tumours: Incidence and time trends in 13 Southern and Eastern European cancer registries. Eur J Cancer 2015; 51: 1444–55.

36. Stekhoven DJ, Buhlmann P. MissForest--non-parametric missing value imputation for mixed-type data. Bioinformatics 2012; 28: 112–8.

37. Lacour B, Guyot-Goubin A, Guissou S, Bellec S, Desandes E, Clavel J. Incidence of childhood cancer in France: National Children Cancer Registries, 2000-2004. Eur J Cancer Prev 2010; 19: 173–81.

38. Kaatsch P, Spix C. German Childhood Cancer Registry - Report 2013/14 (1980-2013). Institute of Medical Biostatistics, Epidemiology and Informatics (IMBEI) at the University Medical Center of the Johannes Gutenberg University Mainz, Germany; 2014.

39. Katanoda K, Shibata A, Matsuda T, Hori M, Nakata K, Narita Y, Ogawa C, Munakata W, Kawai A, Nishimoto H. Childhood, adolescent and young adult cancer incidence in Japan in 2009-2011. Jpn J Clin Oncol 2017; 10.1093/jjco/hyx070: 1-10.

40. United States Department of Health and Human Services CfDCaPaNCI. United States Cancer Statistics: 1999 - 2013 Incidence, WONDER Online Database. 2016.

41. Ellison L, Janz T. Childhood cancer incidence and mortality in Canada: Statistics Canada; 2015.

42. Schmidt LS, Schmiegelow K, Lahteenmaki P, Trager C, Stokland T, Grell K, Gustafson G, Sehested A, Raashou-Nielsen O, Johansen C, Schuz J. Incidence of childhood central nervous system tumors in the Nordic countries. Pediatr Blood Cancer 2011; 56: 65–9.

43. Fairley L, Picton SV, McNally RJ, Bailey S, McCabe MG, Feltbower RG. Incidence and survival of children and young people with central nervous system embryonal tumours in the North of England, 1990-2013. Eur J Cancer 2016; 61: 36–43.

44. Smith MA, Freidlin B, Ries LA, Simon R. Trends in reported incidence of primary malignant brain tumors in children in the United States. J Natl Cancer Inst 1998; 90: 1269–77.

45. Spycher BD, Lupatsch JE, Zwahlen M, Roosli M, Niggli F, Grotzer MA, Rischewski J, Egger M, Kuehni CE, Swiss Pediatric Oncology G, Swiss National Cohort Study G. Background ionizing radiation and the risk of childhood cancer: a census-based nationwide cohort study. Environ Health Perspect 2015; 123: 622–8.

46. Miglioretti DL, Johnson E, Williams A, Greenlee RT, Weinmann S, Solberg LI, Feigelson HS, Roblin D, Flynn MJ, Vanneman N, Smith-Bindman R. The Use of Computed Tomography in Pediatrics and the Associated Radiation Exposure and Estimated Cancer Risk. Jama Pediatrics 2013; 167: 700–7.

47. Spector LG, Pankratz N, Marcotte EL. Genetic and nongenetic risk factors for childhood cancer. Pediatr Clin North Am 2015; 62: 11–25.

48. Spycher BD, Feller M, Roosli M, Ammann RA, Diezi M, Egger M, Kuehni CE. Childhood cancer and residential exposure to highways: a nationwide cohort study. Eur J Epidemiol 2015; 30: 1263–75.

49. Spycher BD, Lupatsch JE, Huss A, Rischewski J, Schindera C, Spoerri A, Vermeulen R, Kuehni CE. Parental occupational exposure to benzene and the risk of childhood cancer: A census-based cohort study. Environ Int 2017; 108: 84–91.

