## Supplemental Figure 1 for "Temporal trends in incidence of childhood cancer in Switzerland, 1985-2014"

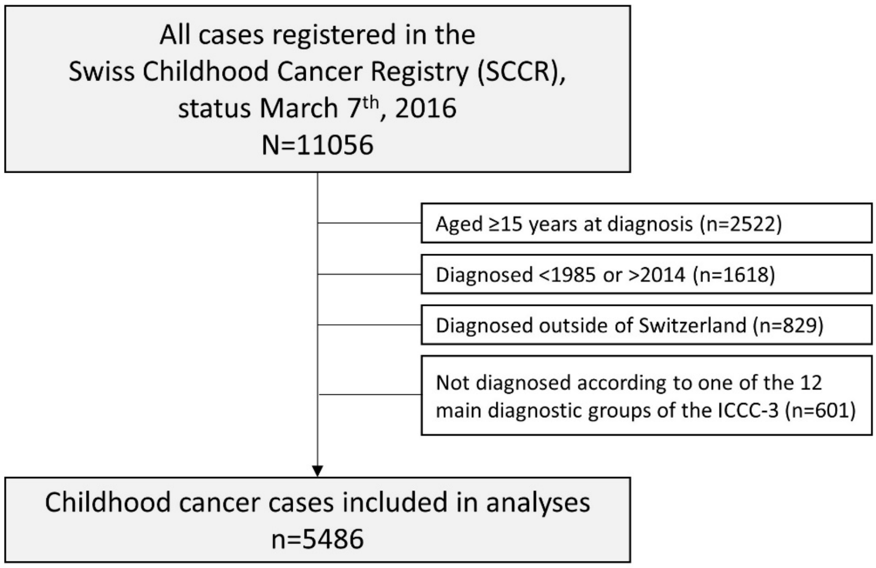


**Supplemental Figure 1.** Flow diagram from the number of registered cases in the SCCR to those eligible for the study.

**Abbreviations**: ICCC-3, International Classification of Childhood Cancer, third edition.
