## Supplemental Table 1 for "Temporal trends in incidence of childhood cancer in Switzerland, 1985-2014"

**Supplemental Table 1a.** Incidence of childhood cancer in Switzerland at age 0-14 years from 2005-2014, by main diagnostic group and subgroup*

| Diagnostic group/subgroup | Average number of cases/year (%) | ASR per million population/year (95% CI)[a] | CR per million population (95% CI)[b] | 1:N children[c] | Male/Female[d] |
|---|---|---|---|---|---|
| *All cancers* | 191.4 (100) | 161.4 (154.2; 168.7) | 2410.6 (2382.8; 2438.4) | 415 | 1.28 |
| I. Leukaemias | 64.3 (33.6) | 54.5 (50.3; 58.8) | 816.0 (799.8; 832.2) | 1225 | 1.57 |
| a. Lymphoid leukaemias | 51.7 (27.0) | 43.9 (40.1; 47.7) | 657.3 (642.8; 671.9) | 1521 | 1.47 |
| b. Acute myeloid leukaemias | 7.8 (4.1) | 6.6 (5.1; 8.1) | 98.4 (92.8; 104.0) | 10163 | 1.89 |
| c. Chronic myeloproliferative diseases | 0.9 (0.5) | 0.7 (0.3; 1.2) | 11.1 (9.2; 13.0) | 90090 | 3.50 |
| d. Myelodysplastic syndrome and other myeloproliferative diseases | 3.2 (1.7) | 2.7 (1.8; 3.6) | 40.3 (36.7; 43.9) | 24814 | 3.00 |
| e. Unspecified and other specified leukaemias | 0.7 (0.4) | 0.6 (0.2; 1.0) | 8.9 (7.2; 10.6) | 112360 | 0.75 |
| II. Lymphomas | 21.2 (11.1) | 17.5 (15.1; 19.8) | 261.1 (251.9; 270.2) | 3830 | 1.75 |
| a. Hodgkin lymphomas | 9.6 (5.0) | 7.8 (6.2; 9.3) | 115.7 (109.6; 121.8) | 8643 | 0.88 |
| b. Non-Hodgkin lymphomas (except Burkitt lymphoma) | 5.7 (3.0) | 4.7 (3.5; 6.0) | 71.0 (66.2; 75.8) | 14085 | 2.80 |
| c. Burkitt lymphoma | 5.4 (2.8) | 4.5 (3.3; 5.7) | 68.0 (63.3; 72.6) | 14706 | 6.71 |
| d. Miscellaneous lymphoreticular neoplasms | 0.5 (0.3) | 0.4 (0.1; 0.8) | 6.4 (5.0; 7.9) | 156250 | 0.25 |
| III. CNS tumours | 43.9 (22.9) | 36.8 (33.4; 40.3) | 553.0 (539.7; 566.4) | 1808 | 1.17 |
| a. Ependymomas and choroid plexus tumour | 4.8 (2.5) | 4.1 (3.0; 5.3) | 61.1 (56.7; 65.6) | 16367 | 1.29 |
| b. Astrocytomas | 18.4 (9.6) | 15.4 (13.1; 17.6) | 231.2 (222.6; 239.8) | 4325 | 1.07 |
| c. Intracranial and intraspinal embryonal tumours | 8.0 (4.2) | 6.8 (5.3; 8.2) | 101.8 (96.1; 107.5) | 9823 | 1.35 |
| d. Other gliomas | 5.1 (2.7) | 4.3 (3.1; 5.5) | 64.9 (60.4; 69.5) | 15408 | 1.32 |
| e. Other specified intracranial and intraspinal neoplasms | 6.7 (3.5) | 5.5 (4.2; 6.8) | 82.8 (77.6; 87.9) | 12077 | 1.16 |
| f. Unspecified intracranial and intraspinal neoplasms | 0.9 (0.5) | 0.8 (0.3; 1.2) | 11.2 (9.3; 13.1) | 89286 | 0.80 |
| IV. Neuroblastoma | 11.8 (6.2) | 10.4 (8.5; 12.2) | 151.7 (144.7; 158.7) | 6592 | 1.11 |
| a. Neuroblastoma and ganglioneuroblastoma | 11.8 (6.2) | 10.4 (8.5; 12.2) | 151.7 (144.7; 158.7) | 6592 | 1.11 |
| b. Other peripheral nervous cell tumour | 0 | | | | |
| V. Retinoblastoma | 4.2 (2.2) | 3.7 (2.6; 4.8) | 54.1 (49.9; 58.3) | 18484 | 0.83 |
| VI. Renal tumours | 9.7 (5.1) | 8.4 (6.7; 10.0) | 124.4 (118.1; 130.7) | 8039 | 0.80 |
| a. Nephroblastoma and other nonepithelial renal tumours | 9.2 (4.8) | 7.9 (6.3; 9.6) | 118.3 (112.1; 124.5) | 8453 | 0.80 |
| b. Renal carcinomas | 0.5 (0.3) | 0.4 (0.1; 0.8) | 6.1 (4.7; 7.5) | 163934 | 0.67 |
| c. Unspecified renal tumours | 0 | | | | |
| VII. Hepatic tumours | 1.5 (0.8) | 1.3 (0.6; 2.0) | 19.1 (16.7; 21.6) | 52356 | 2.00 |
| a. Hepatoblastoma | 1.4 (0.7) | 1.2 (0.6; 1.9) | 18.0 (15.6; 20.4) | 55556 | 1.80 |
| b. Hepatic carcinomas | 0.1 (0.1) | 0.1 (0.0; 0.2) | 1.2 (0.5; 1.8) | 833333 | n.a. |
| c. Unspecified hepatic tumours | 0 | | | | |

**Supplemental Table 1b (continued).** Incidence of childhood cancer in Switzerland at age 0-14 years from 2005-2014, by main diagnostic group and subgroup*

| Diagnostic group/ subgroup | Average number of cases/year (%) | ASR per million population/year (95% CI)[a] | CR per million population (95% CI)[b] | 1:N children[c] | Male/ Female[d] |
|---|---|---|---|---|---|
| VIII. Bone tumours | 7.9 (4.1) | 6.5 (5.0; 7.9) | 97.1 (91.5; 102.7) | 10299 | 0.88 |
| a. Osteosarcomas | 3.8 (2.0) | 3.1 (2.1; 4.1) | 46.6 (42.7; 50.4) | 21459 | 0.90 |
| b. Chondrosarcomas | 0.1 (0.1) | 0.1 (-0.1; 0.2) | 1.2 (0.5; 1.8) | 833333 | n.a. |
| c. Ewing tumour and related sarcomas of bone | 3.9 (2.0) | 3.2 (2.2; 4.2) | 48.2 (44.3; 52.2) | 20747 | 0.95 |
| d. Other specified bone tumours | 0 | | | | |
| e. Unspecified malignant bone tumours | 0.1 (0.1) | 0.1 (0.0; 0.2) | 1.2 (0.5; 1.8) | 833333 | n.a. |
| IX. Soft tissue sarcomas | 13.9 (7.3) | 11.6 (9.7; 13.6) | 173.9 (166.5; 181.4) | 5750 | 1.36 |
| a. Rhabdomyosarcomas | 7.8 (4.1) | 6.6 (5.1; 8.1) | 98.5 (92.9; 104.1) | 10152 | 1.36 |
| b. Fibrosarcomas, peripheral nerve sheath tumours, other fibrous neoplasms | 1.0 (0.5) | 0.8 (0.3; 1.3) | 12.3 (10.3; 14.3) | 81301 | 2.33 |
| c. Kaposi sarcoma | 0 | | | | |
| d. Other specified soft tissue sarcomas | 4.0 (2.1) | 3.3 (2.3; 4.3) | 49.4 (45.4; 53.4) | 20243 | 0.90 |
| e. Unspecified soft tissue sarcomas | 1.1 (0.6) | 0.9 (0.4; 1.5) | 13.8 (11.7; 15.9) | 72464 | 4.50 |
| X. Germ cell tumours | 5.7 (3.0) | 4.8 (3.6; 6.1) | 71.0 (66.2; 75.8) | 14085 | 1.19 |
| a. Intracranial and intraspinal germ cell tumours | 1.5 (0.8) | 1.2 (0.6; 1.9) | 18.5 (16.1; 21.0) | 54054 | 2.00 |
| b. Malignant extracranial and extragonadal germ cell tumours | 1.7 (0.9) | 1.5 (0.8; 2.2) | 21.9 (19.2; 24.5) | 45662 | 0.70 |
| c. Malignant gonadal germ cell tumours | 2.4 (1.3) | 2.0 (1.2; 2.8) | 29.3 (26.3; 32.4) | 34130 | 1.18 |
| d. Gonadal carcinomas | 0 | | | | |
| e. Other and unspecified malignant gonadal tumours | 0.1 (0.1) | 0.1 (0.0; 0.3) | 1.3 (0.6; 1.9) | 769231 | n.a. |
| XI. Epithelial neoplasms & melanomas | 6.9 (3.6) | 5.6 (4.3; 7.0) | 84.0 (78.8; 89.2) | 11905 | 0.60 |
| a. Adrenocortical carcinomas | 0.2 (0.1) | 0.2 (0.0; 0.4) | 2.5 (1.6; 3.4) | 400000 | 1.00 |
| b. Thyroid carcinomas | 1.4 (0.7) | 1.1 (0.5; 1.7) | 16.8 (14.5; 19.1) | 59524 | 0.27 |
| c. Nasopharyngeal carcinomas | 0.2 (0.1) | 0.2 (0.0; 0.4) | 2.4 (1.5; 3.2) | 416667 | 1.00 |
| d. Malignant melanomas | 1.5 (0.8) | 1.2 (0.6; 1.9) | 18.5 (16.0; 20.9) | 54054 | 0.50 |
| e. Skin carcinomas | 0.6 (0.3) | 0.5 (0.1; 0.9) | 7.5 (6.0; 9.1) | 133333 | 2.00 |
| f. Other and unspecified carcinomas | 3.1 (1.6) | 2.5 (1.6; 3.4) | 37.5 (34.1; 41.0) | 26667 | 0.63 |
| XII. Other malignant neoplasms | 0.4 (0.2) | 0.3 (0.0; 0.7) | 5.0 (3.8; 6.3) | 200000 | n.a. |
| a. Other specified malignant tumours | 0.2 (0.1) | 0.2 (0.0; 0.4) | 2.6 (1.7; 3.5) | 384615 | 1.00 |
| b. Other unspecified malignant tumours | 0.1 (0.1) | 0.1 (0.0; 0.3) | 1.3 (0.6; 1.9) | 769231 | n.a. |

**Abbreviations:** ASR, age-standardised incidence rate; CI, confidence interval; CNS, central nervous system; CR, cumulative incidence rate.

* Analyses exclude 98 cases from death certificate notifications because data on specific diagnostic subgroups was not available.

[a] Standardised according to the 2010 European standard population; [b] Cumulative incidence up to the age of 14 years; [c] Number of children affected up to the age of 14 years in Switzerland; [d] male:female ratio.
