## Supplemental Table 2 for "Temporal trends in incidence of childhood cancer in Switzerland, 1985-2014"

**Supplemental Table 2.** Proportions of patients treated in paediatric cancer centres[a] among all children diagnosed with cancer in Switzerland at age 0-14 years from 1985-2014, by diagnostic period

| | 1985-1994 | | | | 1995-2004 | | | | 2005-2014 | | | | |
|---|---|---|---|---|---|---|---|---|---|---|---|---|---|
| | **All cases** | **Paediatric cancer centres[a]** | **%** | **95% CI** | **All cases** | **Paediatric cancer centres[a]** | **%** | **95% CI** | **All cases** | **Paediatric cancer centres[a]** | **%** | **95% CI** | **p-value[b]** |
| *All cancers (N=5486)* | 1673 | 1407 | 84.1 | 82.3; 85.9 | 1880 | 1748 | 93.0 | 91.8; 94.1 | 1933 | 1891 | 97.8 | 97.2; 98.5 | <0.001 |
| *Age at diagnosis* | | | | | | | | | | | | | |
| 0-4 years | 804 | 689 | 85.7 | 83.3; 88.1 | 825 | 791 | 95.9 | 94.5; 97.2 | 830 | 818 | 98.6 | 97.7; 99.4 | <0.001 |
| 5-9 years | 444 | 372 | 83.8 | 80.4; 87.2 | 498 | 469 | 94.2 | 92.1; 96.2 | 509 | 496 | 97.4 | 96.1; 98.8 | <0.001 |
| 10-14 years | 425 | 346 | 81.4 | 77.7; 85.1 | 557 | 488 | 87.6 | 84.9; 90.3 | 594 | 577 | 97.1 | 95.8; 98.5 | <0.001 |
| *ICCC-3 Main diagnostic group* | | | | | | | | | | | | | |
| I. Leukaemias | 569 | 518 | 91.0 | 88.7; 93.4 | 581 | 567 | 97.6 | 96.3; 98.8 | 647 | 644 | 100 | 99.0; 99.9 | <0.001 |
| II. Lymphomas | 201 | 179 | 89.1 | 84.7; 93.4 | 225 | 213 | 94.7 | 91.7; 97.6 | 214 | 211 | 98.6 | 97.0; 99.9 | <0.001 |
| III. CNS tumours | 328 | 234 | 71.3 | 66.4; 76.2 | 429 | 390 | 90.9 | 88.2; 93.6 | 448 | 437 | 97.5 | 96.1; 99.0 | <0.001 |
| IV. Neuroblastoma | 134 | 117 | 87.3 | 81.7; 92.9 | 115 | 113 | 98.3 | 95.9; 99.9 | 119 | 118 | 99 | 97.5; 99.9 | <0.001 |
| V. Retinoblastoma | 46 | 41 | 89.1 | 80.1; 98.1 | 53 | 49 | 92.5 | 85.3; 99.6 | 42 | 40 | 95.2 | 88.8; 99.9 | 0.566 |
| VI. Renal tumours | 102 | 99 | 97.1 | 93.8; 99.9 | 98 | 97 | 99.0 | 97.0; 99.9 | 98 | 98 | 100 | | 0.186 |
| VII. Hepatic tumours | 18 | 11 | 61.1 | 38.6; 83.6 | 27 | 27 | 100 | | 16 | 15 | 94 | 81.9; 99.9 | <0.001 |
| VIII. Bone tumours | 73 | 63 | 86.3 | 78.4; 94.2 | 95 | 91 | 95.8 | 91.8; 99.8 | 79 | 79 | 100 | | <0.001 |
| IX. Soft tissue sarcomas | 111 | 93 | 83.8 | 76.9; 90.6 | 129 | 115 | 89.1 | 83.8; 94.5 | 139 | 136 | 97.8 | 95.4; 99.9 | <0.001 |
| X. Germ cell tumours | 45 | 37 | 82.2 | 71.1; 93.4 | 53 | 51 | 96.2 | 91.1; 99.9 | 57 | 54 | 94.7 | 88.9; 99.9 | 0.026 |
| XI. Epithelial neoplasms & melanomas | 31 | 11 | 35.5 | 18.6; 52.3 | 65 | 30 | 46.2 | 34.0; 58.3 | 69 | 55 | 79.7 | 70.2; 89.2 | <0.001 |
| XII. Other malignant neoplasms | 15 | 4 | 26.7 | 4.3; 49.0 | 10 | 5 | 50.0 | 19.0; 81.0 | 5 | 4 | 80 | 44.9; 99.9 | 0.107 |

**Abbreviations:** CI, confidence interval; CNS, central nervous system; ICCC-3, International Classification of Childhood Cancer, third edition.

[a]Includes the nine clinics of the Swiss Paediatric Oncology Group, [b]p-value derived from Kruskal-Wallis rank sum test for trend, comparing patient numbers treated in paediatric cancer centres between the three time periods.
