## Supplemental Table 3 for "Temporal trends in incidence of childhood cancer in Switzerland, 1985-2014"

**Supplemental Table 3.** Incidence rates of childhood cancer per one million children in Switzerland, other European countries, the USA, Canada, Australia and Korea

| *Country* | | **Switzerland** | | **Piedmont, Italy** | **Spain** | **Sweden** | **France** | **Germany** | **Austria** | **Western Europe** | **USA** | **Canada** | **Australia** | **Korea** |
|---|---|---|---|---|---|---|---|---|---|---|---|---|---|---|
| *Period* | 1985-1994 | 1995-2004 | 2005-2014 | 1967-2011 | 1982-2002 | 1990-1998 | 2000-2004 | 2005-2014 | 2009-2011 | 2001-2010 | 2000-2013 | 2006-2010 | 1997-2006 | 1999-2011 |
| *First author, year of publication* | | Sommer, 2018 | | Isaevska, 2017 | Peris-Bonet, 2010 | Dreifaldt, 2004 | Lacour, 2010 | Kaatsch, 2014 | Karim-Kos, 2016 | Steliarova-Foucher, 2017 | US Cancer Statistics, 2016 | Statistics Canada, 2015 | Baade, 2010 | Park, 2016 |
| All cancers | 143 | 154 | 162 | 157 | 156 | 173 | 157 | 167 | 169 | n.i. | 167 | 161 | 158 | 135 |
| *ICCC-3 Main diagnostic groups* | | | | | | | | | | | | | | |
| I. Leukaemias | 49 | 47 | 55 | 51 | 46 | 51 | 46 | 57 | 54 | 51 | 54 | 52 | 53 | 46 |
| II. Lymphomas | 17 | 18 | 18 | 19 | 18 | 19 | 17 | 16 | 14 | 16 | 17 | 18 | 15 | 13 |
| III. CNS tumours | 28 | 35 | 37 | 37 | 33 | 49 | 36 | 40 | 44 | 38 | 35 | 31 | 36 | 18 |
| IV. Neuroblastoma | 12 | 10 | 11 | 12 | 15 | 11 | 15 | 14 | 12 | 13 | 13 | 13 | 10 | 12 |
| V. Retinoblastoma | 4 | 5 | 4 | 4 | 5 | 5 | 5 | 4 | 5 | 5 | 5 | n.i. | 4 | 5 |
| VI. Renal tumours | 9 | 8 | 8 | 7 | 8 | 10 | 10 | 10 | 9 | 10 | 10 | n.i. | 9 | 6 |
| VII. Hepatic tumours | 2 | 2 | 1 | 2 | 2 | 3 | 1 | 2 | 1 | 2 | 3 | n.i. | 3 | 3 |
| VIII. Bone tumours | 6 | 8 | 7 | 9 | 8 | 5 | 7 | 6 | 10 | 7 | 6 | n.i. | 7 | 7 |
| IX. Soft tissue sarcomas | 9 | 11 | 12 | 9 | 10 | 8 | 10 | 10 | 10 | 10 | 11 | 10 | 8 | 7 |
| X. Germ cell tumours | 4 | 4 | 5 | 4 | 4 | 5 | 6 | 5 | 5 | 5 | 5 | 5 | 6 | 10 |
| XI. Other epithelial neoplasms | 3 | 5 | 6 | 4 | 6 | 4 | 4 | n.i. | 7 | 4 | 6 | n.i. | 7 | 5 |
| XII. Other malignant neoplasms | 1 | 1 | 0.4 | 0.8 | 0.5 | 5 | 0.3 | 0.2 | 0 | 0.3 | 0.7 | n.i. | 0.4 | 3 |

**Abbreviations:** CNS, central nervous system; ICCC-3, International Classification of Childhood Cancer, third edition; n.i., not indicated.
