## Supplemental Table 4 for "Temporal trends in incidence of childhood cancer in Switzerland, 1985-2014"

**Supplemental Table 4a.** Average annual percentage change (95% CI) of childhood cancer in Switzerland, other European countries, the USA, Canada, Australia, Korea and China

| *Country*<br>*Period*<br>*First author, year of publication* | **Switzerland**<br>1985-2014<br>Sommer, 2018 | **Sweden**<br>1960-1998<br>Dreifaldt, 2004 | **Great Britain**<br>1966-2005<br>Kroll, 2012 | **Piedmont, Italy**<br>1976-2011<br>Isaevska, 2017 | **Spain**<br>1983-2002<br>Peris-Bonet, 2010 | **Western Germany**<br>1987-2004<br>Spix, 2008 | **Eastern Germany**<br>1991-2004<br>Spix, 2008 | **Austria**<br>1994-2011<br>Karim-Kos, 2016 | **Ireland**<br>1994-2011<br>National Cancer Registry Ireland, 2014 |
|---|---|---|---|---|---|---|---|---|---|
| All cancers | **0.7 (0.5; 1.0)** | **1.0 (0.8; 1.2)** | **1.0 (0.9; 1.1)** | **1.1 (0.8; 1.5)** | **1.9 (1.5; 2.4)** | **0.8 (0.6; 1.1)** | **2.1 (1.2; 2.9)** | 0.7 (-0.2; 1.7) | Girls:<br>0.8 (-0.2; 1.8)<br>Boys:<br>0.6 (-0.8; 2.0) |
| I. Leukaemias | **0.8 (0.2; 1.4)** | **0.9 (0.4; 1.3)** | **0.7 (0.6; 0.9)** | **0.6 (0.0; 1.2)** | **ALL and AML combined:**<br>**1.6 (0.8; 2.3)** | **0.6 (0.2; 1.1)** | **2.1 (0.7; 3.5)** | 0.3 (-0.8; 1.4) | 0.7 (-1.0; 2.4) |
| II. Lymphomas | 0.0 (-0.9; 1.0) | **1.9 (1.2; 2.6)** | **0.8 (0.6; 1.1)** | **1976-2007:**<br>**1.7 (0.6; 2.7)**<br>2007-2011:<br>-12.2 (-33.2; 15.4) | | **0.9 (0.2; 1.6)** | -0.9 (-3.0; 1.2) | -0.7 (-2.2; 0.8) | -1.3 (-3.8; 1.3) |
| III. CNS tumours | **1985-2002:**<br>**3.0 (1.3; 4.6)**<br>2000-2014:<br>-0.2 (-2.4; 2.1) | **1.5 (1.0; 1.9)** | **1.3 (1.2; 1.5)** | **1.8 (0.9; 2.7)** | **2.2 (1.2; 3.2)** | **1.1 (0.6; 1.6)** | **5.5 (4.6; 7.4)** | **2.9 (0.9; 5.0)** | 1994; 1999:<br>-6.3 (-12.9; 0.7)<br>1999; 2011:<br>2.7 (0.7; 4.7) |
| IV. Neuroblastoma | -0.5 (-1.7; 0.7) | **1.6 (0.8; 2.4)** | **0.6 (0.4; 0.9)** | **1.2 (0.2; 2.1)** | n.i. | 0.1 (-1.6; 1.8) | 2.3 (-0.8; 5.5) | -4.1 (-9.8; 1.6) | n.i. |
| V. Retinoblastoma | -0.9 (-3.0; 1.2) | 0.3 (-1.2; 1.7) | **0.6 (0.2; 1.1)** | 0.7 (-11.1; 14.2) | n.i. | -0.8 (-2.5; 1.0) | n.a. | n.a. | n.i. |
| VI. Renal tumours | -0.3 (-1.5; 0.9) | 0.3 (-0.5; 1.1) | **0.7 (0.4; 1.0)** | 0.2 (-1.1; 1.5) | n.i. | 0.6 (-0.4; 1.6) | 1.8 (-1.5; 5.2) | 0.7 (-2.3; 3.7) | n.i. |
| VII. Hepatic tumours | n.a. | **2.6 (2.0; 3.2)** | **2.5 (1.7; 3.3)** | 3.8 (-13.4; 24.4) | n.i. | 1.5 (-0.9; 3.9) | n.a. | n.a. | n.i. |
| VIII. Bone tumours | 0.2 (-0.9; 1.4) | 0.2 (-0.8; 1.3) | **0.5 (0.2; 0.9)** | 0.0 (-4.1; 4.4) | n.i. | -0.1 (-1.3; 1.0) | 0.6 (-2.9; 4.3) | -0.2 (-3.3; 2.8) | n.i. |
| IX. Soft tissue sarcomas | 1.0 (-0.4; 2.5) | 0.1 (-0.8; 1.0) | **1.6 (1.3; 1.9)** | 0.5 (-0.7; 1.7) | n.i. | **1.3 (0.3; 2.3)** | -0.6 (-3.7; 2.6) | 0.6 (-1.9; 3.0) | n.i. |
| X. Germ cell tumours | 1.6 (-0.2; 3.4) | **1.2 (0.2; 2.2)** | **1.1 (0.6; 1.6)** | 2.4 (-5.9; 11.5) | n.i. | -0.2 (-1.5; 1.1) | -1.1 (-5.6; 3.5) | 2.4 (-2.3; 7.0) | n.i. |
| XI. Epithelial neoplasms & melanomas | **3.8 (1.7; 6.0)** | 0.0 (-1.1; 1.1) | **1.8 (1.3; 2.3)** | 3.9 (-1.4; 9.5) | n.i. | 2.0 (-0.7; 4.8) | n.a. | n.a. | n.i. |
| XII. Other malignant neoplasms | n.a. | **0.4 (0.01; 0.9)** | **2.2 (1.2; 3.2)** | n.a. | n.i. | n.a. | n.a. | n.a. | n.i. |

**Supplemental Table 4b (continued).** Average annual percentage change (95% CI) of childhood cancer in Switzerland, other European countries, the USA, Canada, Australia, Korea and China

| *Country* | **USA** | **USA** | **USA** | **Canada** | **Australia** | **Korea** | **China** |
|---|---|---|---|---|---|---|---|
| *Period* | 1975-2002 | 1992-2004 | 2001-2009 | 1992-2006 | 1997-2006 | 1999-2011 | 2000-2010 |
| *First author, year of publication* | Ward, 2006 | Linabery, 2006 | Siegel, 2014 | Mitra, 2012 | Baade, 2010 | Park, 2016 | Zheng, 2015 |
| All cancers | **0.7*** | 0.4 (-0.1; 0.8) | 0.3 (-0.1; 0.7) | 0.0 (-0.5; 0.4) | **1983-1994: 1.7 (0.9; 2.5)** 1994-2006: -0.1 (-0.7; 0.6) | **2.4 (2.1; 2.7)** | **2.8 (1.1; 4.6)** |
| I. Leukaemias | **0.7*** | 0.7 (-0.1; 1.5) | 0.5 (-0.3; 1.3) | **1992-1999: 2.4 (0.0; 4.9)** 1999-2002: -4.4 (-20.1; 14.2) 2002-2006: -3.0 (-2.6; 9.0) | **0.9 (0.3; 1.5)** | increase‡ | n.i. |
| II. Lymphomas | **-0.5*** | stable‡ | 0.5 (-0.2; 1.3) | 0.0 (-1.4; 1.4) | **0.7 (0.0; 1.3)** | increase‡ | n.i. |
| III. CNS tumours | **1.2*** | -0.1 (-1.1; 1.0) | -0.1 (-1.0; 0.8) | -0.4 (-1.3; 0.5) | **1983-1998: 1.7 (0.6; 2.8)** 1998-2006: -1.8 (-4.5; 1.0) | stable‡ | n.i. |
| IV. Neuroblastoma | 0.5 | -0.6 (-2.9; 1.7) | -1.2 (-3.0; 0.8) | -0.2 (-1.8; 1.5) | 0.2 (-1.1; 1.4) | **5.6*** | n.i. |
| V. Retinoblastoma | 0.5 | 0.3 (-1.5; 2.1) | -0.2 (-1.8; 1.5) | **-2.6 (-4.7; -0.4)** | 0.2 (-1.1; 1.4) | stable‡ | n.i. |
| VI. Renal tumours | 0.4 | -2.1 (-4.6; 0.4) | 0.5 (-0.3; 1.3) | -1.3 (-3.2; 0.7) | 0.4 (-0.7; 1.6) | stable‡ | n.i. |
| VII. Hepatic tumours | **2.0*** | **4.3 (0.2; 8.7)** | 1.7 (-1.7; 5.3) | 1.6 (-0.8; 4.0) | **3.3 (0.8; 5.9)** | stable‡ | n.i. |
| VIII. Bone tumours | 0.3 | 0.2 (-1.4; 1.8) | -0.6 (-1.3; 0.2) | -1.2 (-2.8; 0.5) | 0.3 (-0.8; 1.3) | stable‡ | n.i. |
| IX. Soft tissue sarcomas | **1.0*** | stable‡ | 0.3 (-1.4; 1.0) | -1.4 (-3.6; 0.8) | -0.2 (-1.4; 1.1) | **increase‡** | n.i. |
| X. Germ cell tumours | n.i. | 0.8 (-0.7; 0.3) | 0.7 (-0.5; 2.0) | -0.4 (-2.2; 1.4) | **2.3 (0.9; 3.7)** | stable‡ | n.i. |
| XI. Epithelial neoplasms & melanomas | n.i. | **2.8 (0.5; 5.1)** | **0.8 (0.1; 1.5)** | 2.5 (-0.5; 5.6) | **1983-1994: 4.3 (1.6; 7.0) 1996-2006: -5.7 (-9.1; -2.2)** | **5.6*** | n.i. |
| XII. Other malignant neoplasms | n.i. | n.i. | 0.6 (-2.9; 4.3) | **4.6 (0.1; 9.4)** | n.i. | **-7.4 (-12.2; -2.3)** | n.i. |

**Abbreviations:** ALL, acute lymphoblastic leukaemia; AML, acute myeloid leukaemia; CI, confidence interval; CNS, central nervous system; ICCC-3, International Classification of Childhood Cancer, third edition; n.a., not applicable; n.i., not indicated. * p-value>0.05 (no CI reported); ‡ no total values reported. Bold letters highlight AAPCs whose 95% CIs does not include the null value, or whose p-values<0.05.

**This table does not contain an exhaustive list of all existing reports and studies on incidence trends of childhood cancer. It includes studies and reports who investigate either all childhood cancer diagnoses combined or both childhood leukaemias and CNS tumours.**
